# Gene regulatory networks for compatible versus incompatible grafts identify a role for SlWOX4 during junction formation

**DOI:** 10.1101/2021.02.26.433082

**Authors:** Hannah Thomas, Lisa Van den Broeck, Ryan Spurney, Rosangela Sozzani, Margaret Frank

**Affiliations:** Cornell University, School of Integrative Plant Science, Ithaca, NY 14850, USA; North Carolina State University, Department of Plant and Microbial Biology, Raleigh, NC 27695, USA; North Carolina State University, Department of Electrical and Computer Engineering, Raleigh, NC 27695, USA

## Abstract

Graft incompatibility is a poorly understood phenomenon that presents a serious agricultural challenge. Unlike immediate incompatibility that results in rapid death, delayed incompatibility can take months or even years to manifest, creating a significant economic burden for perennial crop production. To gain insight into the genetic mechanisms underlying this phenomenon, we developed a model system with *Solanum lycopersicum* ‘tomato’ and *Capsicum annuum* ‘pepper’ heterografting, which expresses signs of anatomical junction failure within the first week of grafting. By generating a detailed timeline for junction formation we were able to pinpoint the cellular basis for this delayed incompatibility. Furthermore, we infer gene regulatory networks for compatible self-grafts versus incompatible heterografts based on these key anatomical events, which predict core regulators for grafting. Finally, we delve into the role of vascular development in graft formation and validate SlWOX4 as a regulator for grafting in tomato. Notably, SlWOX4 is the first gene to be functionally implicated in vegetable crop grafting.

## Introduction

Plants have robust systems for self-regeneration following wounding (Savatin et al., 2014; Ikeuchi et al., 2019). Grafting is an ancient agricultural approach that relies on this innate capacity of plants to undergo self-repair. Independent root and shoot systems are surgically joined together, creating a dual plant system that expresses superior traits on either half of the junction. This approach has been strategically adopted in a wide range of species to boost crop productivity and resilience (Mudge et al., 2009; Gaut et al., 2019). Successful grafts are dependent on the formation of the graft junction, a dynamic anatomical connector that unites the rootstock and scion together.

While survival has recently been equated with graft compatibility, the classic definition for compatible combinations states that both non-vascular (cortex/pith, epidermis) and vascular connections must be made between the scion and stock (Proebsting, 1928). Within the Solanaceae, potato, tobacco, and eggplant are routinely grafted with tomato for horticultural purposes (Lee and Oda, 2010; Dawson, 1942). Unlike other Solanaceous plants, Capsicum species (peppers) are only graft compatible with other Capsicum species (Lee and Oda, 2010; Kawaguchi et al., 2008), and tomato and pepper graft combinations have been described as “severely” incompatible (Kawaguchi et al., 2008). The capacity for an incompatible graft to survive for months, or even years in perennial crops, without forming a successful vascular connection is referred to as delayed incompatibility (Argles, 1937). Stunted root and shoot growth, the formation of suckers or adventitious roots, and large, bulging graft junctions are all symptoms of delayed incompatibility (Eames and Cox, 1945; Zarrouk et al., 2006); Copes, 1980). Graft combinations with delayed incompatibility eventually succumb to their mechanical weakness and break at the graft junction, presenting severe challenges for commercial growers (Kawaguchi et al., 2008).

Despite the long history and wide-spread use of grafting, only a small handful of genes are directly implicated in junction formation. These genes are involved in cell proliferation, vascular specification (Asahina et al., 2011; Melnyk et al., 2018; Pitaksaringkarn et al., 2014; Matsuoka et al., 2018; Notaguchi et al., 2020).

Given the essential role of vascular reconnection during graft formation, genes involved in the relatively well-characterized process of cambium-xylem maintenance serve as promising developmental regulators of junction formation. Vascular development in arabidopsis roots is regulated by a dynamic transcription factor network coordinated with hormonal inputs. TRACHEARY ELEMENT DIFFERENTIATION INHIBITORY FACTORs (TDIFs) are produced in the phloem and bind to the PHLOEM INTERCALATED WITH XYLEM (PXY) cambial receptor (Smit et al., 2020; Ito, 2006). Activated PXY is involved in the maintenance of cambial cells by promoting WUSCHEL-RELATED HOMEOBOX 4 (WOX4) and WOX14 (Hirakawa et al., 2010; Etchells and Turner, 2010; Fisher and Turner, 2007; Han et al., 2018; Etchells et al., 2013; Suer et al., 2011). Downstream of WOX14, there are important cambial regulators such as KNOTTED-LIKE FROM ARABIDOPSIS THALIANA (KNAT1) and LOB DOMAIN-CONTAINING PROTEIN 4 (LBD4) (Mele et al., 2003). PXY also represses xylem differentiation factors such as VASCULAR-RELATED NAC-DOMAIN 6 (VND6), VND7, and NAC SECONDARY WALL THICKENING PROMOTING FACTORs (NSTs) via brassinosteroid signaling (Kubo et al., 2005; Zhong et al., 2007; Mitsuda et al., 2007; Kondo et al., 2015; Turco et al., 2019).

In line with the hypothesis that genes involved in xylem-cambial maintenance play a role during junction formation, several core regulators for vascular genesis were identified in recent graft transcriptome studies (Melnyk et al., 2018; Xie et al., 2019). Moreover, these studies uncovered a subset of genes that were asymmetrically expressed either in the scion or the rootstock during graft formation, which lead to an as yet, untested hypothesis that asymmetric expression across the graft interface drives junction formation (Melnyk et al., 2018; Xie et al., 2019).

In this study, we investigate the molecular mechanisms underlying compatible versus incompatible grafts by connecting anatomical processes with predicted regulatory interactions. Through anatomical, biophysical, and genetic characterization, we have established tomato and pepper as a model system for studying graft incompatibility. Only a few studies have employed regulatory networks to identify genes involved in graft formation (Xie et al., 2019). In this study, we utilized Bayesian inference and regression analyses to expand our understanding of species-specific genetic responses which regulate the conserved process of junction formation ((Prill et al., 2010; de Luis Balaguer and Sozzani, 2017; de Luis Balaguer et al., 2017; Clark et al., 2019; Smet et al., 2019). We then identified orthologs of known genetic factors involved in vascular development, which uncovered SlWOX4 as a potential regulator of graft compatibility. In line with this hypothesis, we show that *Slwox4* homografts fail to form xylem bridges across the junction. These functional analyses demonstrate that indeed, *SlWOX4* is essential for vascular reconnection during grafting, and may function as an early indicator of graft failure.

## Results

### Tomato and pepper exhibit delayed incompatibility

To investigate the developmental regulation of graft compatibility, we developed a genetically tractable heterografting system between *Solanum lycopersicum* var. M82 (tomato) and *Capsicum annuum* var. Big Dipper (pepper). In agreement with previous work on tomato and pepper heterografting (Andrews and Marquez, 2010; Kawaguchi et al., 2008), our self-grafted tomato and pepper plants exhibited 100% survival, while hetero-grafted pepper:tomato (scion:stock notation) and tomato:pepper plants showed significantly reduced viability (75% and 37%, respectively; p-value = 8.648e-06; data collected 30 days after grafting; Figure 1M, Supplemental Figure 1A). Furthermore, in contrast to the self-grafted species, the heterografted combinations exhibited reduced foliage, asynchronous stem bulging, and the tomato:pepper grafts displayed severely stunted roots compared to the self-grafts (Figure 1A-D). Fragility and breakage along the junction point is another classic symptom of graft incompatibility (Zarrouk et al., 2006). We performed a bend test to assess whether the biophysical integrity of the pepper and tomato heterografted junctions was significantly reduced (Supplemental Video 1). Only 6% of the self-grafted pepper stems and 0% of the self-grafted tomato stems broke at the junction, while, the majority of the heterografts broke at this position (75% of pepper:tomato stems and 100% of tomato:pepper) (p= 4.405e-07, Fisher’s Exact Test) (Figure 1N, Supplemental Figure 1B).

**Figure 1:**
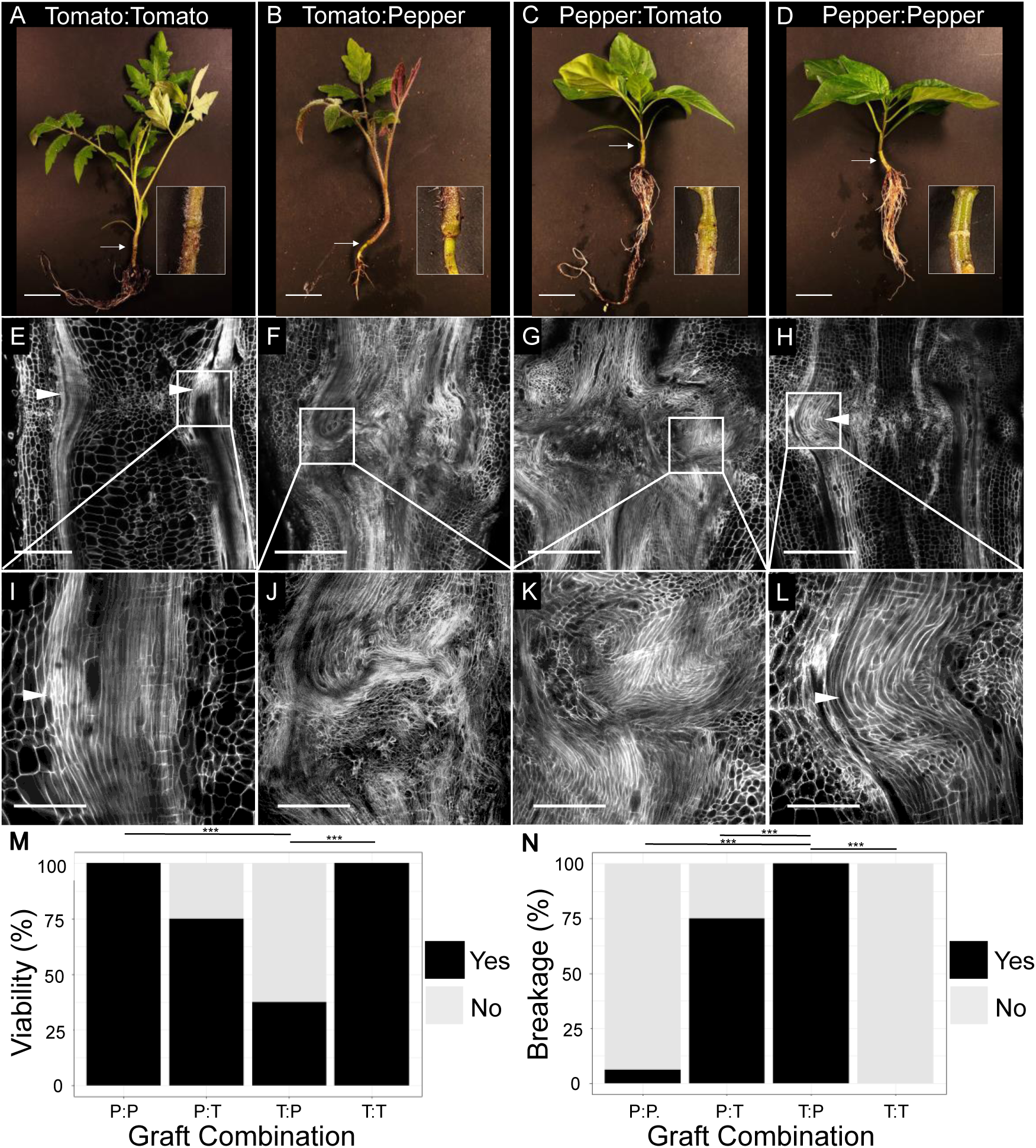
Heterografted tomato and pepper plants show severe vascular patterning defects, reduced viability, and biomechanical failure 30 days after-grafting (DAG). (A-D) Representative images of self-grafted tomato (A), heterografted tomato:pepper (B), pepper:tomato (C), and self-grafted pepper (D) plants taken 30 DAG. White arrows indicate graft junctions. High-resolution confocal imaging of vascular anatomy for self-grafted tomato (E, I), heterografted tomato:pepper (F, J) and pepper:tomato (G, K), and self-grafted pepper (H, L) plants taken at 30 DAG. Tissues were stained with propidium iodide to visualize cell walls, and cleared in methyl salicylate. White arrowheads point to xylem bridges. Heterografts exhibited significantly reduced viability relative to self-grafted plants (M), and higher breakage along the graft site during our bend test (N). “Yes’’ indicates a failure to withstand the bend test, leading to breakage at the graft junction. “No” indicates the stem could withstand the bend test or broke at a secondary location on the stem. For M & N, *** = p-value < 0.001 (Fisher’s Exact Test, contingency tables shown in supplemental figure 1). P:P = pepper:pepper graft, T:T = tomato:tomato graft, P:T = pepper:tomato graft, T:P = tomato:pepper graft. In A-D scale bars = 2 cm, E-H scale bars = 1 cm, I-L scale bars = 400 um.

To identify the cause of graft failure and junction fragility in the heterografts, we inspected the cellular and anatomical detail of the self- and heterografted junctions at 30 DAG (Figure 1E-L, Supplemental Figure 1C-F). Continuous xylem files span the graft junction in the self-grafted tomato and pepper plants, indicating that nutrient and water flow was restored between the scion and stock (Figure 1E, 1H). Our anatomical imaging showed that these new xylem strands were formed toward the periphery of the junction, creating a thickened xylem bridge (Figure 1E, 1H; Mng’omba et al., 2007). Conversely, the heterografts showed an overproliferation of disorganized metaxylem above and below the graft interface (Figure 1F-G, 1J-K). These masses of disconnected xylem files are known as anastomoses and signify a breakdown in the vascular continuity of the stem (Tiedemann, 1989). Despite fully healed epidermal and cortical layers across the junction, all of the heterografted samples failed to form vascular bridges (Figure 1F-G). This data supports a model where heterografted tomato:pepper and pepper:tomato have delayed incompatibility due to failed vascular reconnection.

### Differences between compatible versus incompatible graft anatomy form within the first week of grafting

The formation of functional vascular tissue is crucial for successful grafting. Our heterografts exhibit severe disruptions in vascular strand reconnection. To identify when these vascular phenotypes manifest, we constructed an anatomical timeline for junction formation, comparing self-grafted tomato and pepper with heterografted tomato:pepper and pepper:tomato junctions between 3-6 DAG (Figure 2). We observed parenchymatous callus formation, especially along the stem periphery in all graft combinations (Figure 2). Self-grafted tomatoes exhibited significant callus production at 3 DAG (Figure 2A), and early differentiation of bulbous callus cells into proxylem by 3-4 DAG (Figure 2B). We distinguished these transitioning callus-to-protoxylem cells based on the combination of their isometric shape and characteristic spiral cell wall thickenings (Figure 2A-B; Esau, 1965). The vasculature continued to differentiate 5-6 DAG (Figure 2C-E), which led to elongated xylem strands that connected across the graft junction by 6 DAG (Figure 2D-E).

**Figure 2:**
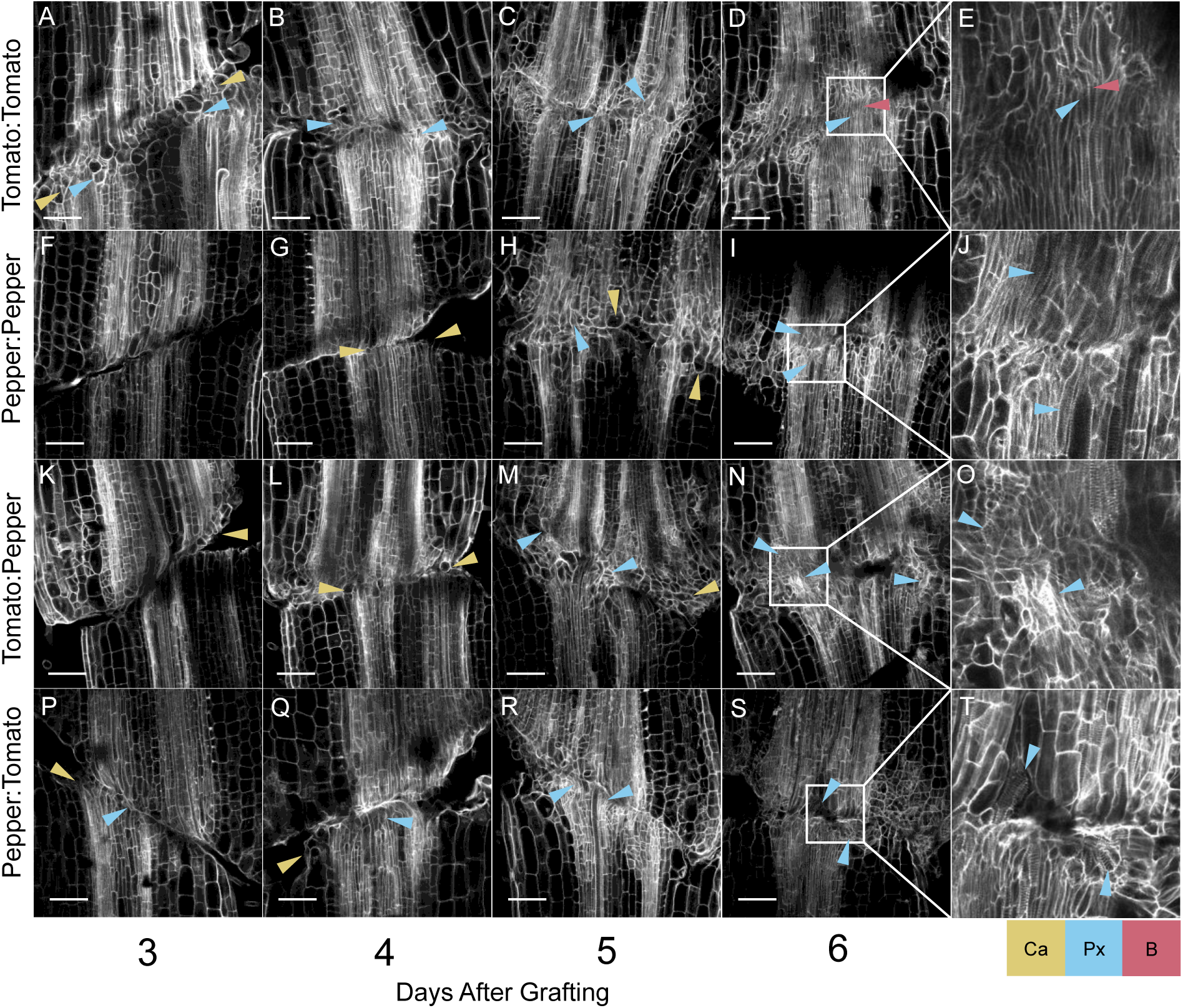
Tomato and pepper heterografts express graft incompatibility within the first week post-grafting. Anatomical timeline for self-grafted tomato (A-E) and pepper (F-J), and heterografted tomato:pepper (K-O) and pepper:tomato (P-T) collected 3-6 days after grafting shows delayed vascular progression and xylem discontinuity in heterograft combinations. Newly formed callus cells are marked with yellow arrowheads, newly formed protoxylem cells are marked with blue arrowheads, and xylem bridges are marked with red arrowheads. The tissue was stained with Propidium Iodide and cleared in methyl salicylate. Scales bars = 200 µm.

In contrast to tomato self-grafts, self-grafted pepper stems showed significant water loss during junction formation. This, in combination with a slower rate of callus formation, increased the fragility of the pepper grafts. While pepper roughly followed the same anatomical stages as tomato, it lagged behind by about 24 hours, potentially due to the increased fragility of the junction. Accordingly, we identified callus cells at 4 DAG (Figure 2G), bulbous callus-protoxylem cells at 5 DAG (Figure 2H), and early signs of vascular maturation by 6 DAG (Figure 2I-J). Much like self-grafted tomato, we observed a considerable amount of callus production in tomato:pepper and pepper:tomato heterografts along the tomato half of the junction at 3 DAG (Figure 2K, 2P). Moreover, we identified protoxylem formation between 3-5 DAG in both heterografts, but again, this was only on the tomato side of the junction (Figure 2M, 2P-S). Thus, while the tomato half of the heterografts exhibited parenchymatous and vascular proliferation, pepper stems remained developmentally stalled during the first 5 DAG, exhibiting no signs of protoxylem differentiation until 6 DAG (Figure 2N-0, 2S-T). Pepper and tomato self-grafts exhibit mild differences (24 hrs) in the temporal development of the junction; however, when heterografted, pepper exhibits a strongly delayed wound response that leads to the discoordination of vascular patterning across the junction. Unlike the self-grafted plants that formed mature vascular connections by 6 DAG (Figure 2D-E, I-J), we did not observe any xylem bridges across the heterograft interface, demonstrating that failed vascular connectivity manifests early in the development of this incompatible combination.

### Molecular networks support distinct hub regulators for self-grafted tomato and pepper

To identify genetic regulators that are essential for proper vascular patterning in the graft junction, we generated temporal gene regulatory networks (GRNs) for graft formation in compatible self-grafts and incompatible heterografts. Using our anatomical timeline, we selected informative sample points that are associated with crucial steps during graft formation: graft adhesion (1 DAG), callus formation (3 DAG), and protoxylem differentiation (5 DAG) (Figure 2).

To generate this molecular timeline, we harvested junctions for RNA-sequencing from self-grafted and heterografted tomato and pepper combinations at 1, 3, and 5 DAG. Using pairwise comparisons amongst all three timepoints, we identified 497, 530, and 536 differentially expressed genes (DEGs: FDR < 0.05) 1, 3, and 5 DAGs (respectively) in the tomato:tomato self-grafts. (Supplemental Dataset 1). Next, we applied a selection method using all graft-related GO-terms (372) based on our observations from the anatomical timeline and published studies on grafting (Figure 2; Supplemental Dataset 2; Supplemental Figure 2; Melnyk et al., 2018; Xie et al., 2019). To construct GRNs, we included DEGs that overlapped with the graft-related GO-terms (Supplemental Dataset 2, Supplemental Figure 2), as well as all differentially expressed transcription factors (TFs) (Supplemental Figure 3). We identified 168 graft-related DEGs and 63 TFs that we used to perform network inference and predict causal regulations with a Dynamic Bayesian Network (DBN) algorithm (Spurney et al., 2020; de Luis Balaguer et al., 2017; de Luis Balaguer and Sozzani, 2017). To ensure high accuracy, we inferred three networks for each time point and combined the regulatory interactions, visualizing time point-specific, and common regulations (Figure 3, Supplemental Dataset 3).

**Figure 3.**
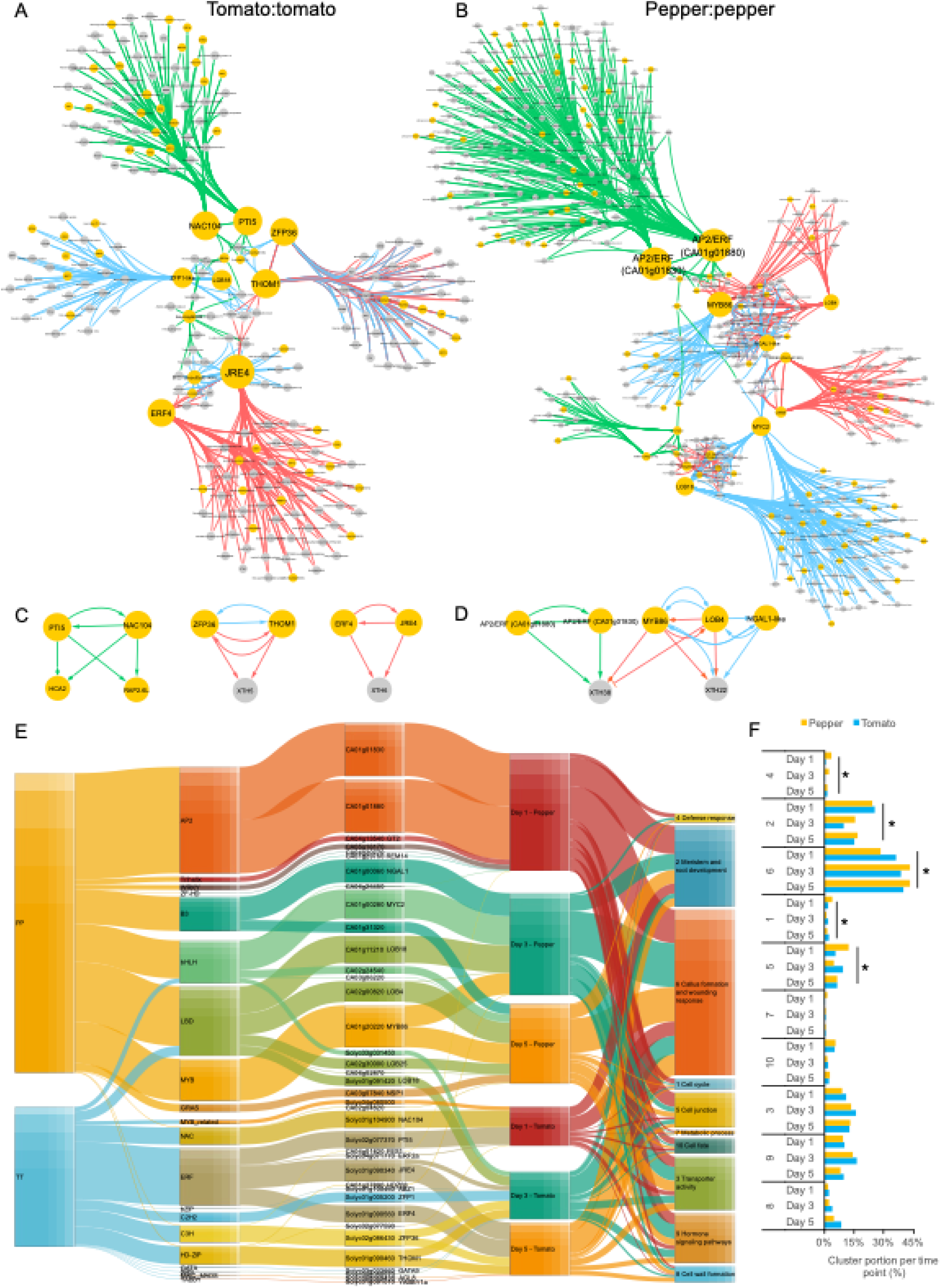
Time-specific modules and their major regulators identified in tomato:tomato and pepper:pepper self-graft gene regulatory networks. (A-B) Causal relations were predicted with a dynamic Bayesian network approach between differentially expressed transcription factors and DEGs associated with Gene Ontology categories related to grafting for the (A) tomato:tomato self-graft and (B) pepper:pepper self-graft. Green, blue and red arrows represent regulations at 1 DAG, 3 DAG, and 5 DAG, respectively. Yellow and grey nodes represent transcription factors and non-transcription factors, respectively. (C-D) Highlighted inferred interactions in the main text from the tomato:tomato (C) and pepper:pepper (D) networks. (E) Sankey diagram visualizing inferred gene regulatory interactions from the tomato:tomato and pepper:pepper networks. The width of the connections between each vertical block represents the number of genes (from left to right): contained within each graft combination network, within each TF family, downstream of the major hub, expressed at a specific time point, and that fall into a specific GO-cluster. All TFs that have an outdegree > 0 are included. (F) Percentage of the downstream target genes associated with each GO-cluster per time point. * = p-value < 0.05 (Fisher exact test).

Within the tomato:tomato network, we identified three subnetworks or modules for each of the time points, as well as a module common for two time points (3 and 5 DAG) (Figure 3A). The early time point module (1 DAG) contains 85 genes that are predominantly regulated by two TFs: an ortholog to NAC104 (Solyc01g104900), which is known to negatively regulate cell death during vascular formation, and an ERF/AP2 protein PTO INTERACTING 5 (PTI5, Solyc02g077370) (Sari et al., 2019; Gu et al., 2002; Gupta et al., 2020). Within this early temporal module, we predict that NAC104 and PTI5 are controlling the expression of tomato orthologs for two arabidopsis genes that are functionally implicated in grafting: RAP2.6L (Solyc12g042210) and HCA2 (Solyc06g071480) (Figure 3C; Miyashima et al., 2019; Asahina et al., 2011). In agreement with the anatomical observations at 3 DAG, when callus cells start to form (Figure 2A), our network predicts two major hubs for cell proliferation: LBD18 (Solyc01g091420) and THOM1 (Solyc01g090460). Notably, LBD18 functions in callus specification and maintenance, and THOM1 marks meristematic cells, both of which are crucial developmental processes during junction formation (Ikeuchi et al., 2017; Meissner and Theres, 1995). Moreover, we infer that THOM1 regulates an XYLOGLUCAN ENDOTRANSGLUCOSYLASE/HYDROLASE (XTH) gene (Solyc07g052980) (Figure 3C). In Arabidopsis, XTH genes, including XTH19 and XTH20, have been shown to function in the proliferation of the pith during tissue regeneration (Pitaksaringkarn et al., 2014). Furthermore, our network infers an additional hub-XTH interaction during the late time point module (5 DAG), where ETHYLENE RESPONSE FACTOR (ERF4; Solyc01g090560), and JASMONATE RESPONSIVE ERF (JRE4; Solyc01g090340) co-regulate a downstream XTH (Solyc11g065600) (Nakayasu et al., 2018) (Figure 3C). Overall, this analysis uncovers newly predicted regulators that control downstream genes with established roles in tissue regeneration and junction formation.

Our anatomical timeline for self-grafted pepper predicts delayed development in junction formation relative to self-grafted tomato. To investigate how molecular networks for graft formation are shifted between these species, we constructed a comparative GRN for pepper. We identified 1318, 683, and 540 DEGs at 1, 3, and 5 DAG, respectively for the self-grafted pepper dataset (Supplemental Dataset 4). We selected graft-related DEGs and TFs following the same guidelines that we applied to self-grafted tomato gene selection (Supplemental Figure 3, Supplemental Dataset 3; Spurney et al., 2020; de Luis Balaguer et al., 2017). This network analysis included 105 TFs and 333 graft-related DEGs (Supplemental Figure 3, Figure 3B). Congruent with our tomato:tomato network analysis, we identified time-specific modules within the pepper:pepper network as well as TFs that are involved in regulating multiple time points. Within the early time point module (1 DAG), we identify two ERF TFs (CA01g01830, CA01g01880) as central regulators in the network (Figure 3B). This contrasts with the tomato:tomato network, where ERFs play a key role at later stages of junction formation (Figure 3A). Furthermore, we identify MYC2 (CA01g00280), involved in jasmonate signaling, LBD18 (CA01g11210), NGAL1-like (CA01g00060), and MYB86 (CA01g20220) as major regulators of junction formation 3 DAG (Dombrecht et al., 2007; Soyano et al., 2008; Lee et al., 2009; Fan et al., 2012; Patzlaff et al., 2003). Notably, MYB86, which has previously been associated with lignification during xylem formation (Patzlaff et al., 2003), functions as a hub at both 3 and 5 DAG. Gene clusters that are downstream of MYB86 are associated with xylem formation, including numerous peroxidase genes, NAC-related TFs, and HOMEOBOX LEUCINE ZIPPER-14 (HD-ZIP 14) (Kajala et al., 2020; Marjamaa et al., 2009). We predicted additional hubs 5 DAG, including LBD4 (CA02g00820) and LBD25 (CA02g30000), which were recently implicated in the related process of haustorium formation during plant parasitism (Jhu et al., 2021; Melnyk, 2017). Finally, we identified multiple interactions where hub genes (LBD4, MYB86, NGAL1-like, and two ERFs (CA01g01880 and CA01g01830)) converged to regulate two XTH genes: XTH22 (CA07g00520) and XTH38 (CA11g08350) (Figure 3D). These hub-XTH modules are similar to the multi-gene regulatory modules that we found in the self-grafted tomato GRN (Figure 3C). Despite similarities in these downstream targets, we uncover distinct hub genes between our self-grafted tomato and pepper GRNs.

Next, we compared the regulations of the pepper and tomato self-grafts to 1-align the networks with our anatomical timeline, 2-identify the specific transcriptional regulations involved in the differential progression of junction formation, and 3-contrast gene networks for self-grafted pepper and tomato. To this end, we used a Sankey diagram, which allows for the comparison and visualization of the number of target genes across different samples, time points, and TF families (Figure 3E). In the diagram, the species, TF family, TFs with at least one downstream target, and the ten selected GO clusters related to grafting, are connected based on the number of their downstream target genes. As expected, AP2/ERF TFs, which were prominent hubs in both pepper and tomato GRNs, are uncovered as key regulators for self-grafting in both species in the Sankey diagram (Figure 3E). Interestingly, the two hub AP2/ERFs in pepper are predicted to play a key role solely at 1 DAG, while tomato ERFs are major regulators at all time points (Figure 3E). Such differential identification of TF families across the time points is also observed for bHLH, LBD, NAC, C3H, and HD-ZIPs. The GO-clusters most closely associated with each time point include: cell cycle, meristem/root development, defense response, and cell fate at 1 DAG, transporter activity and hormone-related signaling pathways at 3 DAG, and cell wall formation at 5 DAG (Figure 3F). Although these trends are similar for both species, we observed a 15% increase in genes associated with callus formation and wounding response (Cluster 6) in self-grafted pepper between 1 and 3 DAG, while tomatoes have strong gene membership starting at 1 DAG (Figure 3E). Delayed activation of cluster 6 in pepper provides further molecular support for our anatomical timeline (Figure 2, Figure 3F).

The Sankey diagram indicates that the difference in the developmental timing of junction formation between self-grafted pepper and tomato originates from the delayed induction of key TF families, such as the LBD family, and/or the absence of other key families at specific time points, for example, the AP2/ERF and NAC families. Additionally, the regulation of DEGs associated with callus formation and wounding response is delayed in the pepper self-grafts. Thus, our network analysis informs us on the molecular underpinnings for the developmental delay in pepper graft formation.

### Incompatible heterografts display severely perturbed genetic regulation

As shown in the network analysis, tomato and pepper self-grafts utilize distinct pathways to heal following grafting. Because of this, we hypothesized that the inability of tomato and pepper heterografts to form vascular connections could be due to misaligned genetic processes required for vascular differentiation across the junction. To fully explore the disruptions of the genetic regulation between compatible and incompatible grafts, we utilized multiple bioinformatic approaches.

First, to identify shared genetic components, we compared DEGs from self- and heterograft plants, which uncovered 185 shared tomato genes and 401 shared pepper genes (Supplemental Figure 4, Supplemental Dataset 5). To identify causal relationships between TFs and downstream graft-related genes for these large groups of heterografted DEGs, we applied a random forest regression tree approach and generated a Sankey diagram (Supplemental Dataset 6, Supplemental Figure 3, Supplemental Figure 5; Clark et al., 2019). Within this dataset, a considerable number of orthologs for graft-related TFs from arabidopsis were identified in the networks, but rarely in both reciprocal graft combinations, highlighting the fact that the incompatible grafts are disrupted in genetically distinct ways (Supplemental Dataset 6). To shed light on whether the orthologs of these known graft-related genes have similar roles in both species, the expression of all tomato and pepper orthologs for nine functionally characterized, grafting- and vasculature-related genes from arabidopsis (VND6, VND7, WOX4, CVP2, PXY, HCA2, RAP2.6L, ALF4, and ANAC071) were plotted from the self-grafted and heterografted datasets. Notably, these plots show highly perturbed expression in the heterografts, compared to the self-grafts (Figure 5C, Supplemental Figure 6; Asahina et al., 2011; Yamaguchi et al., 2010; Melnyk et al., 2018; Sugimoto et al., 2010; Matsuoka et al., 2018; Pitaksaringkarn et al., 2014; Ji et al., 2010; DiDonato et al., 2004; Smit et al., 2020). To identify additional candidates with spatially or temporally dynamic expression patterns in the heterografts, we used a modified Shannon entropy (MSE) analysis that uncovered 34 transcription factors, nine of which were previously identified in graft co-expression networks (Supplemental Figure 7-8, Supplemental Dataset 7; Xie et al., 2019). We constructed gene regulatory networks using a subset of these TFs, and identified downstream targets that have roles in graft formation, for example we found a bHLH TF that regulates ANAC071 (Supplemental Figure 8C, Supplemental Dataset 7; Asahina et al., 2011).

To investigate how regulatory interactions are altered during incompatible graft formation, we overlaid the connectivity of the tomato:pepper and pepper:tomato heterografts onto our previously constructed self-graft networks (Figure 3A-B, Figure 4, and Supplemental Figure 9). We found that many of the self-grafted hubs showed dramatic changes in outdegree (i.e. -the number of outgoing edges from the hub) for the heterograft networks (Figure 4). For example, LBD18, a callus-related gene that acts as a central hub 3 DAG has high levels of connectivity within self-grafted tomato (outdegree = 38; Figure 4A) and pepper (outdegree = 91; Figure 4D) (Supplemental Figure 9). However, we found greatly reduced connectivity for SlLBD18 in the pepper:tomato graft (outdegree = 2; Figure 4B) and for CaLBD18 in the tomato:pepper graft (outdegree = 2; Figure 4F) (Supplemental Figure 9). Additionally, we predicted THOM1, a meristematic marker, as a major co-regulator of self-grafted tomato 3 and 5 DAG (Figure 4A); however, in the heterografts its regulatory connections have been shifted solely to 5 DAG (Figure 4B-C). This shift in THOM1 regulation is congruent with a model where delayed specification of the vascular meristem (the cambium) is associated with disorganized vascular patterning and delayed incompatibility in the heterografts (Figures 1-2).

**Figure 4.**
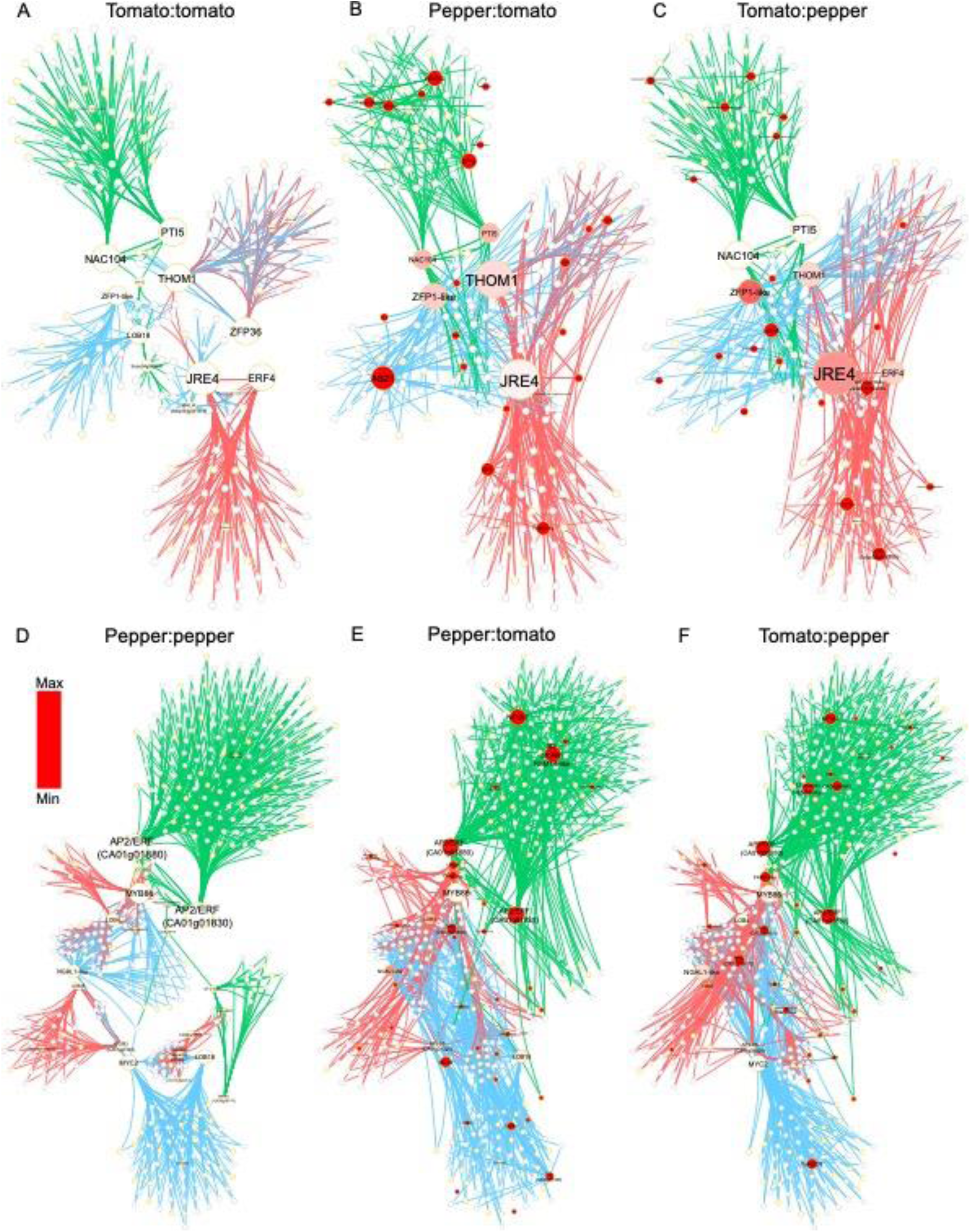
Altered and disrupted regulatory connections in the heterografts. (A-C) The variation in outdegree for the pepper:tomato (B) and tomato:pepper (C) network compared to the tomato:tomato self-graft network (A) is shown. (D-E) Similarly, the variation in outdegree for the pepper:tomato (E) and tomato:pepper (F) network compared to the tomato:tomato self-graft network (D) is shown. Green, blue and red arrows represent regulations at 1 DAG, 3 DAG, and 5 DAG, respectively. Nodes are colored with different shades of red according to the absolute magnitude of their variation in outdegree compared to the self-graft. Yellow and grey bordered nodes represent transcription factors and non-transcription factors, respectively.

**Figure 5.**
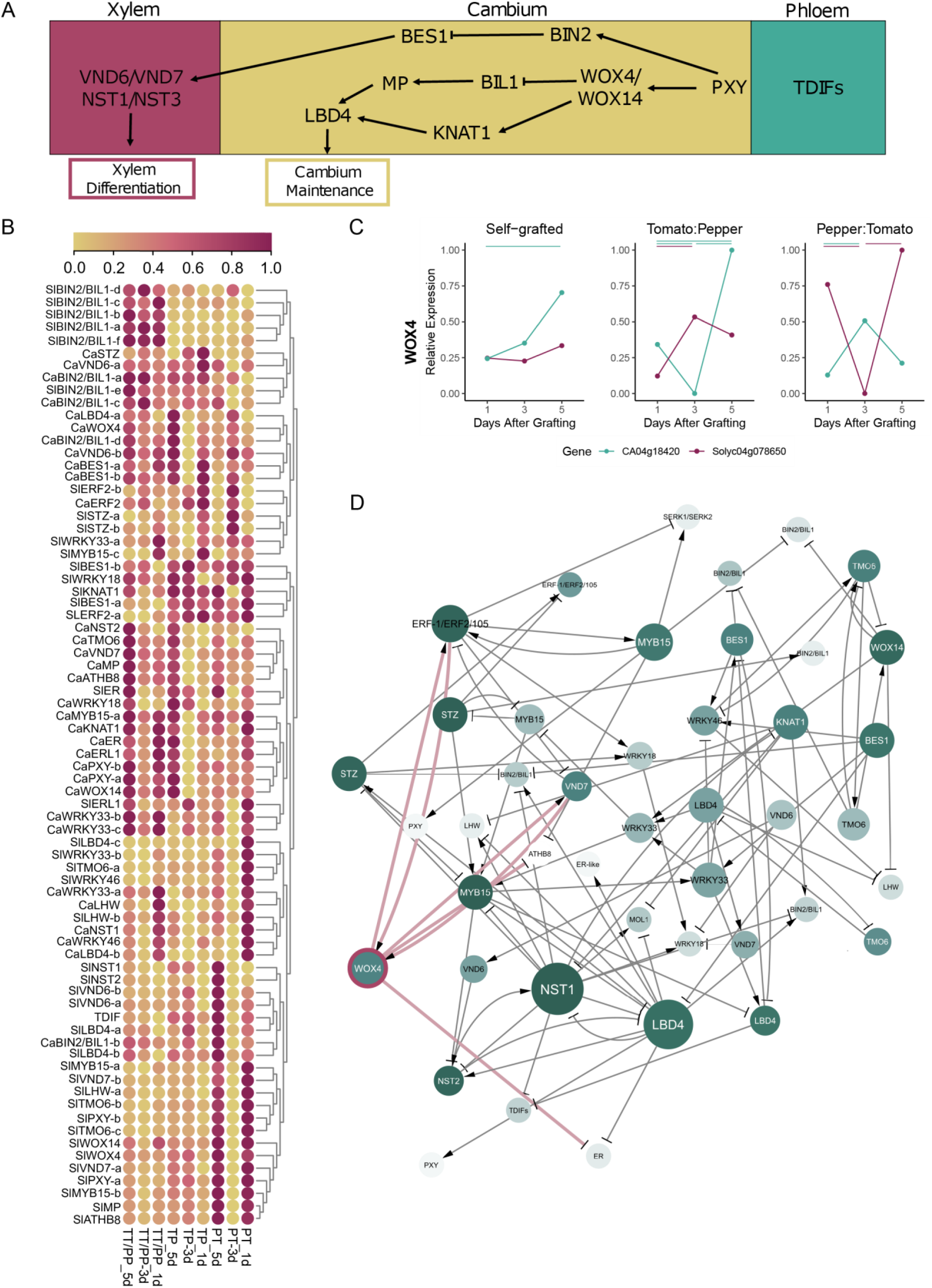
Genes involved in cambium-xylem maintenance are disrupted in heterografted plants. (A) Schematic overview of a network of transcription factors, with WOX4 and WOX14 as central TFs, that underlies the balance between cambial maintenance and xylem differentiation. (B) Scaled expression of the tomato and pepper orthologs of genes involved in cambium-xylem maintenance, depicted in A. (C) Expression pattern of SlWOX4 and CaWOX4 in self-grafted and heterografted plants. Bars show significant differential expression between time points (FDR < 0.05 and log_2_ fold change > 1 or < −1). Blue and red bars signify significant differential expression between pepper and tomato time points, respectively. (D) Inferred regulatory interactions between the tomato orthologs of the genes depicted in A. Nodes are colored according to the magnitude of their variation in edge connections between the heterografts. Node size represents the number of outgoing interactions. WOX4 and its edges are highlighted in pink.

### SlWOX4 is a new regulator for xylem reconnection during graft formation

Our anatomical and molecular analyses demonstrate that graft incompatibility between tomato and pepper is strongly associated with shifts in gene regulatory interactions that lead to failed vascular reconnection. To understand the cause of vascular disruption, we focused our analysis on tomato and pepper orthologs of arabidopsis genes that are involved in specifying and maintaining vascular development (Figure 5A-B). Many of these orthologs exhibited altered expression dynamics between the compatible self-grafts and incompatible heterografts (Supplemental Figure 10). Using a regression approach, we inferred the regulatory interactions between these genes (Figure 5D). Notably, our network inference predicts that VNDs and NSTs, which are both involved in xylem differentiation, are regulated by WOX4 in both the self-grafts and heterografts (Figure 5C, Supplemental Fig 10). We decided to examine SlWOX4 (Solyc04g078650) and CaWOX4 (CA04g18420) in more detail. While these WOX4 orthologs exhibit patterns of gradually elevated expression in self-grafted plants, their expression becomes disrupted and chaotic in the heterografts (Figure 5B-C). These results, in combination with our observation that xylem files fail to form in the incompatible heterografts, led us to the hypothesis that WOX4 may serve a crucial function during graft formation, and disruption of this gene may lead to graft-incompatibility.

To test this hypothesis, we obtained a CRISPR-Cas9 knockout for *Slwox4*. Notably, WOX4 is a critical player in procambial specification and exhibits scion-dominant expression during graft junction formation in arabidopsis (Melnyk et al., 2018; Ji et al., 2010; Etchells et al., 2013). We observed that the overall morphology and anatomy of *Slwox4* was relatively similar to our wild type control, albeit the plants were slightly smaller (Figure 6A-D). To test whether the disrupted expression pattern of cambium-xylem maintenance genes seen with heterografts is associated with incompatibility, we made self- and heterograft combinations between *Slwox4* mutants and wild type controls and evaluated survival as well as anatomical connectivity within the junction. We did not observe a statistically significant difference in the survival rate of *Slwox4* mutant versus wild type grafts at 30 DAG (Supplemental Figure 11). However, while viability was not impacted, we discovered that the self-grafted *Slwox4* junctions exhibited failed xylem connectivity and thus are anatomically incompatible (Figure 6F). Similar to the pepper and tomato heterografts, *Slwox4* self-grafts developed over proliferating and disorganized xylem masses on either side of the junction (Figure 6F). In contrast, when *Slwox4* is only on one side of the junction (i.e. in the *Slwox4*:WT and WT:*Slwox4* heterografts), the grafts formed mature xylem connections that spanned the junction and thus did not exhibit graft incompatibility (Figure 6G-H). These results demonstrate that WOX4 is required in at least one half of the graft junction to maintain the cambial cell population; however, it does not matter which half. From these experiments, we conclude that WOX4 plays a crucial role in xylem reconnection during junction formation.

**Figure 6.**
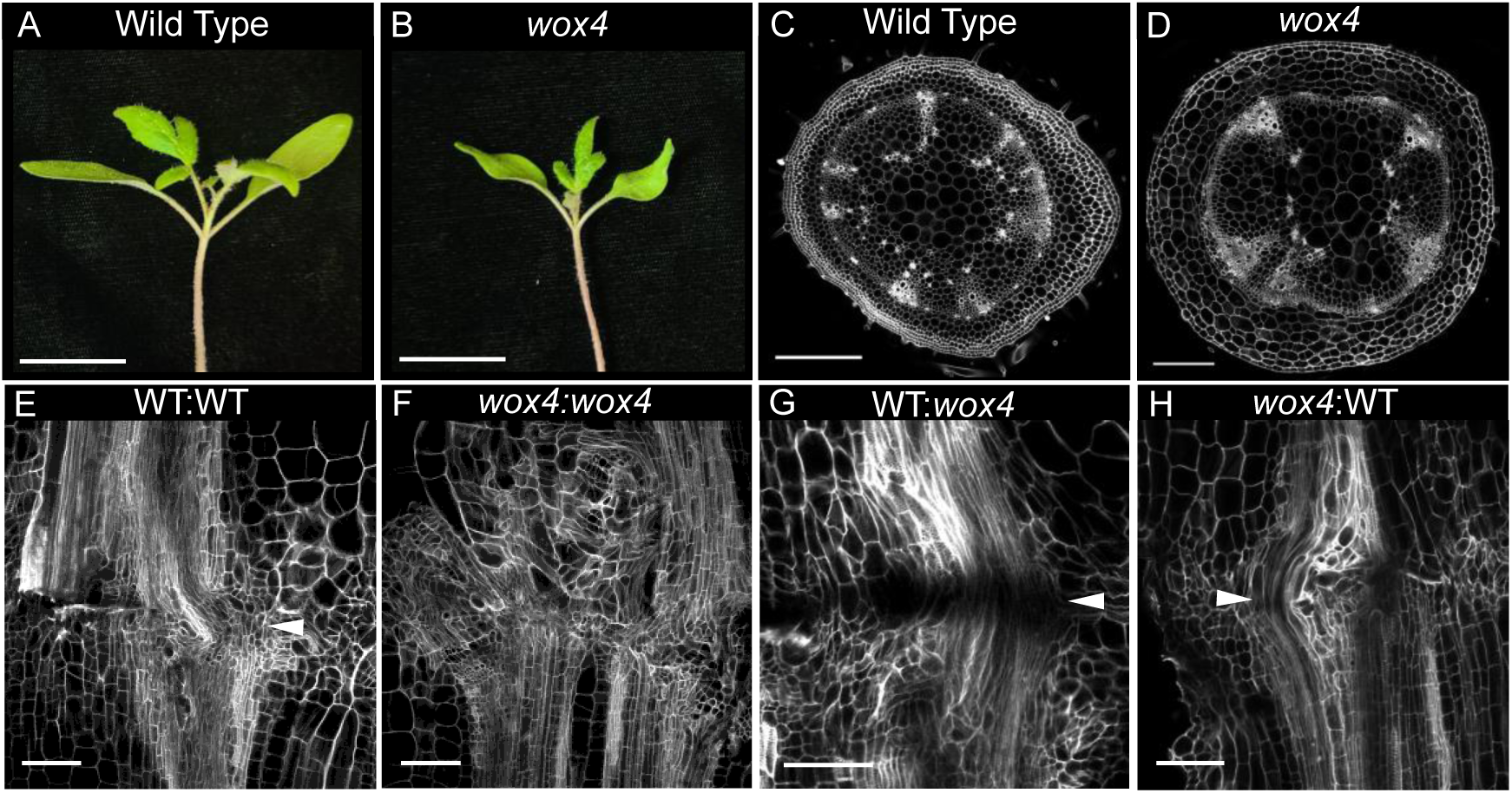
Self-grafted *Slwox4* mutants fail to form xylem bridges, and thus exhibit graft-incompatibility. Representative selection of wild type (A) and *Slwox4* (B) seedlings 3-weeks after imbibition. Representative cross-sections sampled at similar points along the stem: 1 cm above the graft junction, under the first leaf node of wild type (C) and *Slwox4* (D). Representative images of self-grafted wild type (WT) (E), self-grafted Slwox4 (F), WT:Slwox4 (G), and Slwox4:WT (H) 30-days after grafting. Xylem bridges are marked with white arrows. Tissues in C-H were stained with Propidium Iodide and cleared in methyl salicylate. A-B scale bars = 3 cm, C-D scale bars = 500 um, E-H scale bars = 200 um.

## Discussion

The formation of a compatible graft involves the distinct anatomical processes of both non-vascular and vascular healing. While we found that both compatible and incompatible grafts achieved non-vascular healing within one-week post-grafting, our incompatible heterografts failed to form vascular reconnections, even when examined as late as 30 days after grafting (Figures 1-2). However, these incompatible heterografts can survive for several months post-grafting and thus exhibit delayed incompatibility due to failed vascular coordination within the graft junction.

Despite the widespread applications of grafting for agricultural crop improvement, only 7 genes have been directly implicated in graft formation; the majority of which were discovered in arabidopsis (Notaguchi et al., 2020; Pitaksaringkarn et al., 2014; Asahina et al., 2011; Melnyk et al., 2018, 2015). Transcriptomic characterization of junction formation has helped elucidate both temporal and rootstock-scion specific molecular patterns that are associated with graft formation (Melnyk et al., 2018, 2015; Notaguchi et al., 2020; Xie et al., 2019; Pitaksaringkarn et al., 2014; Asahina et al., 2011). However, very few of these studies focus on horticulturally-relevant crops, and none of these studies connect detailed anatomical events with causal network analysis. By constructing our own anatomical timeline (Figure 2) and corresponding, temporal transcriptomic dataset (Figure 3), we synthesize a molecularly informed model for the developmental progression of junction formation (Figure 7). In this model, we predict genetic hubs at 1 DAG that are associated with wound responses, including defense-related genes (PTI5), programmed cell death (NAC104), and ethylene signaling (ERFs). At later developmental stages, 3 and 5 DAG, we identify hubs involved in callus production (LBD18), meristematic activity (THOM1, LBD4, LBD25), and hormonal signaling (AP2/ERFs, MYC2, JRE4, ERF4). Despite genetic diversity in these regulators between tomato and pepper, we show that our hub genes converge on the regulation of functionally related targets that are essential for grafting in arabidopsis (for example, the XTH regulatory modules; Figure 3 C-D).

**Figure 7.**
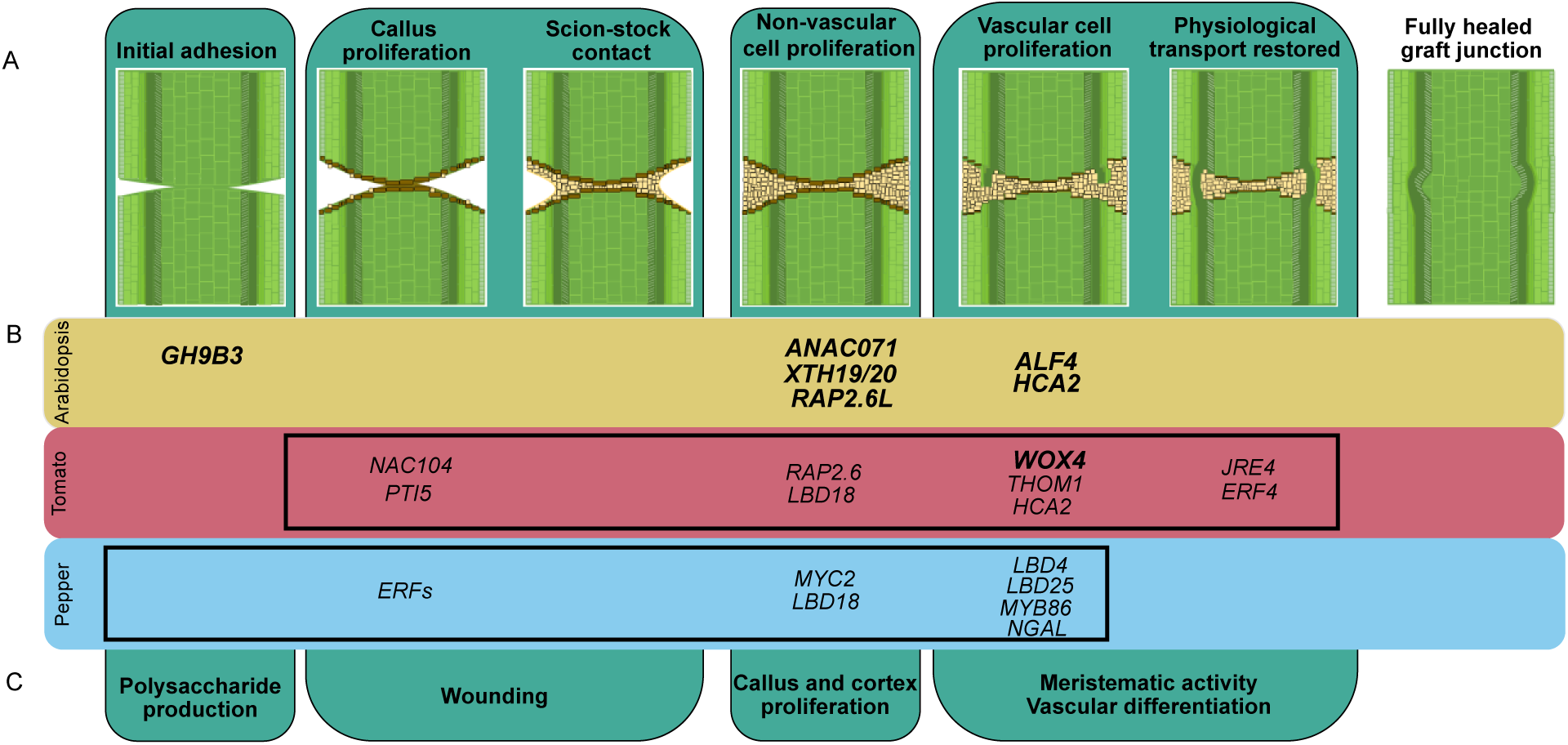
Network hubs predict new and conserved regulators for anatomical reconnection during junction formation. The anatomical timeline conserved throughout graftable plants includes initial adhesion, callus proliferation, scion-stock contact, non-vascular cell proliferation, vascular cell proliferation, and restored physiological transport through reconnected phloem and xylem strands (A). There are seven functionally characterized genes involved in graft junction formation in arabidopsis. We have identified 16 candidate genes for graft junction formation in tomato and pepper, many of which are described for the first time as graft-related, and one of which is the first functionally validated gene involved in vegetable crop graft formation (B). Despite the genetic diversity amongst arabidopsis, tomato, and pepper, all involved genes are associated with core anatomical steps along the graft junction timeline (C). The black boxes in B specify the processes captured in the anatomical timelines for tomato and pepper (Figure 2). Functionally validated genes involved in grafting are bolded.

Previous studies have focused on understanding cell-to-cell interactions in the graft junction, with the aim of identifying graft-specific genetic factors that are independent of wound responses (Melnyk et al., 2018; Xie et al., 2019). We designed our study to investigate the involvement of wound-induced tissue regeneration during junction formation, and thus this work inherently uncovers genetic hubs that were not previously considered to play a role in grafting. These hubs have, however, been implicated in the related process of haustorium formation in parasitic plants, and thus provide molecular support for the connection between graft junctions and haustoria (Melnyk et al., 2018; Xie et al., 2019; Jhu et al., 2021). The fact that we identify diverse regulatory hubs between our species, supports a model in which grafting is not controlled by a genetically conserved process that is evolutionarily programmed into plant genomes. Rather, it is a human invention that draws on the innate capacity for plant regeneration following wounding. In this light, it is logical that the specific genetic regulators for grafting are diverse, while activation of core biological processes related to wound response and regeneration is largely conserved across species.

Because tomato/pepper heterografts fail to form coordinated xylem connections across the graft junction, we looked for orthologs of transcription factors that are involved in cambium-xylem maintenance in arabidopsis (Figure 5A-B). We were able to identify numerous Solanaceae orthologs with disrupted expression patterns in the incompatible heterografts relative to self-grafted tomato and pepper (Figure 5B). By investigating the interconnectivity of these TFs, we identified WOX4 as a central regulator for vascular regeneration during junction formation (Figure 5D). While this role for WOX4 in grafting is logical, given its role in procambial maintenance, the translation of known vascular networks into the identification of genes that are essential for grafting has been challenging (Hirakawa et al., 2010; Etchells et al., 2013; Ji et al., 2010). Our discovery provides a new tool for disrupting graft formation at the crucial stage of xylem reconnection. Despite the apparent disruption of xylem patterning in self-grafted Sl*wox4* junctions, ungrafted *Slwox4* mutants form organized vascular strands, which is likely the result of SlWOX4/SlWOX14 functional redundancy, as was previously demonstrated in arabidopsis (Etchells et al., 2013). Interestingly, we found that heterografted WT:*Slwox4* and *Slwox4*:WT plants form mature xylem bridges across the junction, demonstrating that the requirement of SlWOX4 expression is not directionally specific. This result is somewhat surprising given previous work showing that WOX4 exhibits scion-dominant expression (Melnyk et al., 2018). One possibility is that SlWOX4 is a mobile factor, much like the related WOX family member WUSCHEL, and thus can be expressed on either side of the junction, and still function in the scion (Yadav et al., 2011; Daum et al., 2014).

A long-standing question in the field of grafting, asks whether new vascular bridges develop through the specification of cambium or differentiate directly from callus (McCully, 1983, Roberts, 1949; Tiedemann, 1989; Crafts, 1934). Our discovery of SlWOX4 as an essential regulator of junction formation, indicates that indeed, cambial specification precedes vascular differentiation. The self-grafted *Slwox4* mutants mimic incompatible graft formation, demonstrating an essential role for cambial specification in graft compatibility. Future work delving deeper into cambial patterning within the junction will help resolve how compatible grafts are determined.

## Acknowledgments

We are grateful to Zachary Lippman, Anat Hendelman, and Gina Robitaille for providing CRISPR-Cas9 *wox4* mutant lines. We would also like to thank Adrienne Roeder and Blake Meyers for their helpful suggestions on the manuscript, Jocelyn Rose and Iben Sørensen for assistance with Auromine O staining, and Jennifer Rice for her help developing the pepper/tomato grafting system. This work was supported by Frank Lab startup funds from Cornell University College of Agriculture and Life Sciences. M.H.F was supported by the National Science Foundation (NSF) (CAREER IOS-1942437). H.R.T. was supported by a United States Department of Agriculture National institute of Food and Agriculture (USDA-NIFA) Predoctoral Fellowship (2020-67011-31882); R.S. was supported by the National Science Foundation (NSF) (CAREER MCB-1453130) and the Foundation for Food and Agriculture Research (FFAR). Imaging data was acquired through the Cornell Institute of Biotechnology’s Imaging Facility, with NIH (S10OD018516) funding for the shared Zeiss LSM880 confocal/multiphoton microscope.

## Author contributions

H.R.T, M.H.F., L.V.d.B., and R.S. conceived and designed the study. H.R.T. and M.H.F. gathered experimental data. H.R.T. and L.V.d.B analyzed the data. R.J.S. analyzed the RNAseq data. L.V.d.B. performed network inference and MSE. H.R.T., M.H.F., L.V.d.B., R.J.S, and R.S. wrote the manuscript.

## Methods

### Plant Materials and Growth Conditions

To trigger germination, *Capsicum annuum* (pepper) and *Solanum lycopersicum* (tomato) seeds were treated with 50% bleach for 30 seconds and then rinsed five times with sterile dI-water. Tomato seeds were germinated on wet paper towels in Phytotrays (Sigma-Aldrich) that were placed in the dark for 72 hours, transferred to the light for 72 hours, and then transplanted into LM-111 soil. Pepper seeds were immediately planted 1 cm deep into LM-111 soil. Tomato and pepper seedlings were grown in climate controlled chambers set to 23 C with 16:8 day/night light cycles (500-800 umol/m2/sec).

### Plant growth conditions and grafting

*Capsicum anuum* (var. Big Dipper) seeds were grown as described above. Seven days later *Solanum lycopersicum* (Var. M82) seeds were grown as described above. Twenty-one day old pepper seedlings and 14 day old tomato seedlings, which have the same diameter of stem, were joined with a slant or wedge graft on the internode between the cotyledons and first leaf (Kubota et al., 2008). Grafts were performed in each of the following combinations: tomato:tomato, pepper:pepper, tomato:pepper, and pepper:tomato. Grafts were held together with 1.5 mm silicon-top grafting clips (Johnny’s Selected Seeds, Albion, ME, USA). Grafted plants were generously watered, covered with plastic domes, and placed in the dark for 3 days. On day 4, plants were returned to light (500-800 µmol/m2/sec).

#### Graft compatibility 30 DAG

Fifty *Capsicum anuum* (var. Big Dipper) and 50 *Solanum lycopersicum* (Var. M82) seeds were grown as described above and slant grafted (Kubota et al., 2008). Plastic domes were vented 7 DAG and removed 14 DAG. 16 biological replicates were collected 30 DAG. The junctions were hand-cut longitudinally, and one half was stained with propidium iodide (PI), while the other half was stained with Auramine O (details below). Additional images of propidium iodide stained tissue found in Supplemental Figure 12.

#### Anatomical timeline for graft junction formation

One-hundred-eighty *Solanum lycopersicum* (var. M82) and 180 *Capsicum anuum* (var. Big Dipper) were grown and grafted as described above. Nine biological replicates were collected for each graft combination, 3-6 days after grafting. Stems were fixed in Formalin-Alcohol-Aceitic Acid (FAA), stained with PI, and cleared in methyl salicylate (as described below). Additional images of propidium iodide stained tissue can be found in Supplemental Figure 13.

### CRISPR-Cas9 *Slwox4* targeting

CRISPR-Cas9 gRNA selection and cloning for targeting SlWOX4 was performed by the Lippman lab, as described in previous publications (Kwon et al., 2020; Brooks et al., 2014; Soyk et al., 2019). A binary vector containing two gRNAs targeting SlWOX4 (Solyc04g078650): CR-WOX4-gRNA1-TTGCAACCAAGTGTAAGTGA and CR-WOX4-gRNA2-ATCAAAAGGAGGAGTAACAA were introduced with *Agrobacterium tumefaciens*-mediated transformation into an indeterminate (Sp+) tomato cultivar M82 at the Boyce Thompson Institute Center for Plant Biotechnology Research (Van Eck et al., 2019). First generation transgenic lines were transplanted and genotyped with locus-specific primers (CR-WOX4-conf_F TGGGATCATCATCAGGAAGC and CR-WOX4-conf_R TTAGGAGGGCTATTGCTACTTTCA) as described previously (Soyk et al., 2019).

#### Mutant grafting with *Slwox4*

Indeterminate (Sp+) M82 was used for our wild type control. Fifty *Slwox4* seedlings and 50 wild type seedlings were grown as described above and slant grafted in the following combinations: *wox4*:*wox4,* wild-type:wild-type, *wox4*:wild-type, and wild-type:*wox4*. Plastic domes were vented 7 DAG and removed 14 DAG. Sixteen biological replicates were collected 30 DAG. Additional images of PI stained tissue can be found in Supplemental Figure 14.

### Staining and confocal imaging for graft junction anatomical analyses

#### Tissue collection

Graft junctions were harvested by cutting approximately 1 cm above and below the cut site. Tissue was placed into tissue cassettes (Sakura Finetek USA, Inc. 4117-01), and immediately transferred into ice-cold FAA (10% Formaldehyde, 5% acetic acid, 50% ethyl-alcohol) fixative, and infiltrated under a vacuum for 2-4 hours. The Issue was moved to fresh FAA and stored at 4C overnight. The following day, tissue was moved through an ethanol dehydration series, followed by a rehydration series.

#### Propidium Iodide

After fixing in FAA, and dehydrating and rehydrating tissue, the samples were stained with 20 ug/ml propidium iodide for 1 hour and rinsed with phosphate buffered saline. Tissue was then dehydrated again in the dark, and gradually transferred into methyl salicylate clearing agent. Finally, the tissue was cleared in 100% methyl salicylate at 4 C for 2 weeks. Fully cleared graft junctions were imaged on a Zeiss LSM880 Confocal Microscope using an Argon Laser 514 nm beam.

#### Auramine O

After fixing in FAA, and dehydrating and rehydrating, tissue was stained with 0.01% Auramine O in 0.05M Tris-HCl ph 7.2 for 15 minutes. The tissue was rinsed with water and immediately imaged on a Leica M205 fluorescent dissecting microscope using an EL6000 Mercury Metal Halide light source.

### Bend Test

Graft junction integrity was tested using manual bending. Each stem portion was held 1-2 cm away from the graft site. Even pressure was applied to bend the stem at a 45 degree angle. Stems that broke at the graft junction were marked as broken, stems that did not break or broke at a different point of the stem were considered not broken. All graft combinations: tomato:tomato (n=16), pepper:pepper (n=16), tomato:pepper (n=3), and pepper:tomato (n=12) were tested.

### Imaging

Grafted plants were imaged using a Samsung 12-megapixel wide-angle camera. Seedlings were imaged using Panasonic LUMIX GX85 Mirrorless Camera with a 12-32 mm lens.

### Statistical analysis of grafted plants

Statistical significance of survival and stem integrity was calculated using Fisher’s Exact Test. Pairwise comparisons were conducted using Fisher’s Exact Test. Significance was determined as p < 0.05.

### Construction and Sequencing of RNA-seq libraries

Library construction: 50 *Solanum lycopersicum* (Var. M82) and 50 *Capsicum anuum* (var. Big Dipper) seedlings were grown as described above. Seedlings were wedge grafted. Graft junctions, consisting of 1 cm from the scion and 1 cm from the stock, were harvested between 8-10 PM at 1-day, 3-days, and 5-days post-grafting. Junctions were immediately flash frozen in liquid nitrogen, with 1 junction harvested per biological replicate and 4 biological replicates per time point. RNA was extracted using TRIzol reagent (Thermo Fisher Scientific, Walthm, MA USA). The purified RNA was treated with DNAseI (Thermo Fisher Scientific, Walthm, MA USA), and quantified and quality checked on a DeNovix DS-11 (DeNovix, Willmington, DE) spectrophotometer. RNA-seq libraries were constructed using 2.5 µg of total RNA per sample. Briefly, mRNA sequencing libraries were constructed by isolating mRNA with the NEBNext® Poly(A) mRNA Magnetic Isolation Module (New England Bioloabs, Ipswich, MA USA), followed directly by the NEBNext® Ultra™ Directional RNA Library Prep Kit for Illumina® using NEBNext® Multiplex Oligos for Illumina®. Six libraries were pooled per lane and run as a single-end sequencing run with 101 cycles on an Illumina HiSeq 2500 at the University of Delaware Sequencing and Genotyping Center. All sequencing data are available on GEO at https://www.ncbi.nlm.nih.gov/geo/query/acc.cgi?acc=GSE167482 (GSE167482, access token ufahgmimlzgljyb)

### Bioinformatic analyses

To analyze the time course RNAseq, reads of each sample were mapped against both tomato and pepper reference genomes (Supplemental Dataset 8, Supplemental Figure 15). Averaging across biological replicates and experimental time points, 92.59% and 82.38% reads of the tomato:tomato and pepper:pepper graft junctions were mapped to the tomato and pepper reference genomes, respectively. Mapping the heterografts resulted as expected in ∼50% alignment and a small percentage of reads mapped to the incorrect species. To increase the accuracy of the alignment of the heterograft reads, we performed a concatenation of both reference genomes, in essence, treating the heterografts as hybrids, which resulted in an 87.80% and 87.05% average alignment for the tomato:pepper and pepper:tomato heterografts, respectively. To further explore variability, groupings, and outliers within the time course datasets, we performed a principal component analysis (PCA) that clustered the samples based on the similarity. The four biological replicates clustered close together in PC’s 1 and 2 of the PCA (Supplemental Figure 16). Moreover, in the PCA built using the tomato gene set, the self-grafted samples clustered together compared to the heterografts. In contrast, the PCA built with the pepper gene set showed that the samples clustered according to the time of harvest on the second principal component. Overall, the PCA verified that replicates from the same sample have similar profiles and hinted toward differences between the pepper and tomato response to grafting.

The TuxNet interface was used to perform gene expression analysis and infer GRNs (Spurney et al., 2020). The following genomes were used when running TuxNet: Pepper genome cvCM334 and Tomato genome Solanum lycopersicum cv Heinz (gene version ITAG3.2) (Kim et al., 2014; Tomato Genome Consortium, 2012). TuxNet specifically uses the following softwares: Preprocessing: ea-utils fastq-mcf (Aronesty, 2013), Alignment: hisat2 (Kim et al., 2015), and Differential expression analysis: Cufflinks (Trapnell et al., 2012).

TuxNet also includes an algorithm (TuxOP) for DEG selection using FC and FDR values. Specifically, an FDR threshold of 0.05 and log_2_ FC threshold of 2 was used. DEGs were assigned to a timepoint based on upregulation. For each dataset, up- and down-regulated DEGs were selected from each pairwise comparison: 1 DAG vs. 3 DAG, 1 DAG vs. 5 DAG, and 3 DAG vs. 5 DAG, which captured all temporally regulated DEGs (Supplemental Dataset 1,4,5). To infer a gene regulatory network (GRN), differentially expressed genes (DEGs) associated with one of 372 manually selected GO-terms (Supplemental Dataset 2) as well as all differentially expressed TFs were identified for each of the samples (Supplemental Figure 3). 3951 tomato genes and 4375 pepper genes in the entire tomato and pepper genome, respectively, were associated with one of the 372 graft-related GO-terms (Supplemental Dataset 2, Supplemental Figure 3). To identify the GO-terms associated with each gene, the Gene Ontology of tomato and pepper were downloaded from dicots PLAZA 4.0 (Van Bel et al., 2018). For inferring GRNs for the self-grafts, a dynamic Bayesian network (DBN)-based inference algorithm (GENIST) was used within the TuxNet interface with a time lapse of 0 (de Luis Balaguer et al., 2017). Only putative TF-encoding genes were considered as source nodes that could regulate the expression of other DEGs. Specifically, for each time point for both self-grafts a network was generated using the selected DEGs at that time point and the average expression values from the entire time course. As such three networks were inferred for each self-graft. Finally, we combined the 1, 3, 5 DAG networks by taking the union of the three GENIST output files. The regulatory interactions between the same set of DEGs were inferred within the heterograft networks by using the average expression values from the entire time course of the heterograft for a dynamic Bayesian network approach.

Similarly, one inference was performed for each time point, after which the output of the three inference rounds were unionized in Cytoscape. The networks from the self-grafts and heterografts were compared in Cytoscape through the DyNet application (Goenawan et al., 2016). Specifically, each heterograft network was compared with the self-graft by mapping the variation of outdegree onto the node color.

For the heterograft samples, a random forest approach (RTP-STAR) within the TuxNet interface was used for network inference (Spurney et al., 2020). Similar to the self-grafts, separate networks were generated at each time point for both heterografts by using the selected DEGs and the expression values of all the replicates. Ten iterations were performed in total and the average expression values for the time course were used to determine the sign of the predicted regulations. For each of the two heterograft samples, six networks were generated: one for each time point and species genome. We combined the 1, 3, 5 DAG networks by taking the union of the three RTP-STAR output files. A total of four GRNs were generated, two for each heterograft sample, one for the tomato genes and one for the pepper genes. TuxNet is available at https://github.com/rspurney/TuxNet and video tutorials regarding installation, analysis, and network inference are freely available at https://rspurney.github.io/TuxNet/. All networks were visualized in Cytoscape® 3.8.0 (Shannon et al., 2003). Sankey diagrams were generated in R with the package *networkD3* (Allaire et al., 2017; R Core Team, 2020). The heatmap for Figure 5 was generated using TBtools (Chen et al., 2020), and supplemental heatmaps and plots were generated in R using ggplot2 (Wickham, 2011). For the heterografts, we visualized TFs that were shown to have a major regulatory role (i.e., TFs that regulate more than 25 targets) and selected in total 59 TFs which accounted for >60% of all inferred interactions. To find orthologs across species, we used uniprot (UniProt Consortium, 2019), PANTHER (Mi et al., 2013), and generated custom orthogroupings for *Capsicum annuum*, *Arabidopsis thaliana*, and *Solanum lycopersicum* using the default settings for OrthoFinder (Emms and Kelly, 2015).

Prior to comparative analyses of gene expression values, including the PCA and MSE analysis, between the self-graft aligned to their respective genomes and the heterografts aligned to the concatenated genome, the FPKM values were normalized against the self-graft 1 DAG replicate 1. The PCA was performed in R using the *prcomp* function from the *stats* package. R-code used to perform the MSE is available at https://github.com/LisaVdB/MSE. To perform the semantic clustering of the 382 selected GO-terms, the R package *GOSemSim* was used to compute semantic similarity (Yu, 2020; Yu et al., 2010). The computed similarity matrix was clustered into 10 clusters (optimal number of clusters identified with elbow plot from the within-clusters sum of squares) using k-means clustering.

---

**Supplemental Figure 1: Statistical contingency tables for graft survival and bend test and xylem visualization in graft junction** (Supports Figure 1). (A) Fisher’s exact test for the portion of surviving grafts and (B) the portion of stems that broke at the graft junction during the bend test. The differences amongst graft groups was significantly different in both experiments. (C-F) Representative fluorescent images through hand sections of graft junctions showing xylem bridge formation of self-grafted tomato (C), heterografted tomato:pepper (D) pepper:tomato (E), and self-grafted pepper (F) harvested 30-days after grafting. Lignified cells were stained with Auramine O to xylem profiles. Dashed lines indicate the original graft site. C-F scale bars = 250 uM.

**Supplemental Figure 2 - Clustering of the 372 selected GO terms based on semantic similarity** (Supports Figure 3). A total of 10 GO clusters were identified that include different biological processes related to each other. The percent of overlapping gene membership across GO terms is visualized on a color scale from navy-to-light blue, representing 0-100% overlap.

**Supplemental Figure 3 - Selection of differentially expressed genes (DEGs) associated with grafting for network inference** (Supports Figure 3). For each tissue and species, three sets of genes were compared with Venn diagrams: 1) DEGs from the RNAseq analysis, 2) all known transcription factors for tomato and pepper, and 3) genes related to the 372 selected GO-terms. The number of genes within the overlap between set 1 and 2 and the set 2 and 3 are listed in the left bottom corner of each tissue/species combination and are used for network inference.

**Supplemental Figure 4 - Common differentially expressed genes (DEGs) between the self-grafts and heterografts** (Supports Figure 4). The 185 and 401 overlapping between self-graft and heterografts in tomato and pepper correspond to 80% and 92% of the total DEGs of the self-grafts and to 14% and 26% of the total DEGs of the heterografts, respectively. Genes from the heterograft datasets were selected based on the overlap between temporal DEGs and our selected GO categories.

**Supplemental Figure 5 - Sankey diagram visualizing inferred gene regulatory interactions from the tomato:pepper networks** (Supports Figure 4). The width of the connections between each vertical block represents the number of genes (from left to right): contained within each graft combination network, within each TF family, downstream of the major hub, expressed at a specific time point, and that fall into a specific GO-cluster. All TFs that have an outdegree > 25 are included.

**Supplemental Figure 6 - Graft-specific genes from arabidopsis are disrupted during tomato and pepper hetereografting** (Supports Figure 4). Tomato and pepper orthologs of known genes required for grafting (ANC071, RAP2.6L, HCA2, ALF4), vascular cambium patterning (PXY), protophloem patterning (CVP2), and proto- and metaxylem patterning (VND6, VND7) shown over time in all graft combinations. Expression was normalized across all three genomes and scaled between 0 and 1. Bars show significant differential expression between time points (FDR < 0.05 and log_2_ fold change > 1 or < −1). Blue and red bars signify significant differential expression between pepper and tomato time points, respectively.

**Supplemental Figure 7 - Heatmap of MSE-selected genes** (Supports Figure 4). The heatmap shows the min/max rescaled FPKM values of the 2215 genes selected with modified Shannon entropy (MSE). These 2215 are further subdivided based on their expression pattern in eight groups (from left to right): genes that showed the same dynamical pattern across the self-grafts and heterografts (*All_SamePattern*), genes specifically induced in the heterograft samples (*Heterograft*), genes specifically induced in the self-grafts (*Homograft*), genes that showed an opposite dynamical pattern in the two heterografts (*OppositePattern_heterograft*), genes specifically induced in the scion of pepper (*Scion_specific_pep*), genes specifically induced in the scion of tomato (*Scion_specific_tom*), genes specifically induced in the stock of pepper (*Stock_specific_pep*), genes specifically induced in the stock of tomato (*Stock_specific_tom*).

**Supplemental Figure 8 - Expression pattern of 37 selected TFs** (Supports Figure 4). (A-B) The min/max rescaled expression values of 37 tomato (A) or pepper (B) TFs in common between the modified Shannon entropy (MSE) analysis and GRN inference is shown. These TFs belong to one of eight groups genes that showed the same dynamical pattern across the self-grafts and heterografts (*All_SamePattern*), genes that showed an opposite dynamical pattern in the two heterografts (*Opposite_heterograft*), and genes specifically induced in the heterograft samples (*Heterograft*), the self-grafts (*Homograft*), the scion (*Scion*), or the stock (*Stock*). (C) Transcriptional regulations of three major TFs extracted from the heterograft GRNs that regulate more than 25 downstream targets in the pepper:tomato and tomato:pepper networks, including the graft-related ANAC071.

**Supplemental Figure 9 - Variation in outdegree in the self-grafts and heterografts** (Supports Figure 4). (A-B) The outdegree is plotted of TFs with an outdegree > 2 for the pepper (A) and tomato genes (B).

**Supplemental Figure 10 - WOX4 regulates xylem differentiation genes, VND6/7 and NST1/2, in self-grafts and heterografts** (Supports Figure 5). (A-C) The rewiring of 45 tomato genes in the tomato:tomato (A), tomato:pepper (B) and pepper:tomato (B) network is shown. (D-F) Similarly, the rewiring of 32 pepper genes in the pepper:pepper (D), tomato:pepper (E) and pepper:tomato (F) network is shown. Nodes are colored with different shades of red according to the magnitude of their variation in edge connections. The node and edges of WOX4 are highlighted with a blue border and bolded black arrows, respectively.

**Supplemental Figure 11 - *Slwox4* mutant seedlings do not display decreased viability 30 DAG** (Supports Figure 6). The percentage of surviving plants 30 DAG. Fully wilted plants were considered dead. n=18, * = p-value < 0.05 (Fisher’s Exact Test).

**Supplemental Figure 12: Heterografted tomato and pepper plants show severe vascular patterning defects 30 days after-grafting (DAG)** (Supports Figure 1). Representative images of self-grafted tomato (A-C), self-grafted pepper (D-F), heterografted tomato:pepper (G), heterografted pepper:tomato (H) taken 30 DAG. Figure 12A, 12D, 12G, and 12H are seen at higher magnification in Figure 1A, 1D, 1B, and 1C respectively. The tissue was stained with Propidium Iodide and cleared in methyl salicylate. Scales bars true across all images and = 800 µm.

**Supplemental Figure 13: Tomato and pepper heterografts express graft incompatibility within the first week post-grafting, continued** (Supports Figure 2). (A-I) Anatomical timeline for self-grafted tomato. Self-grafted tomato at 3 DAG (A-B), 4 DAG (C-D), 5 DAG (E), and 6 DAG (F-I). (J-O) Anatomical timeline for self-grafted pepper. Self-grafted pepper at 4 DAG (J-K), 5 DAG (L-M), and 6 DAG (N-O). (P-Z) Anatomical timeline for heterografted pepper:tomato. Pepper:tomato at 3 DAG (P-S), 4 DAG (T-U), 5 DAG (V-X), and 6 DAG (Y-Z). (AA-AK) Anatomical timeline for heterografted tomato:pepper. Tomato:pepper at 3 DAG (AA-AB), 4 DAG (AC-AF), 5 DAG (AG-AH), and 6 DAG (AI-AK). The tissue was stained with Propidium Iodide and cleared in methyl salicylate. Scales bars true across all images and = 800 µm.

**Supplemental Figure 14 - Self-grafted Slwox4 mutants fail to form xylem bridges, and thus exhibit graft-incompatibility, continued** (Supports Figure 6). Representative images of self-grafted wild type (WT) (A-C), self-grafted Slwox4 (D-F), WT:Slwox4 (G-I), and Slwox4:WT (J-L) 30-days after grafting. Figure 14A, 14E, 14I, and 12K are seen at higher magnification in Figure 6E, 6F, 6G, and 6H respectively. The tissue was stained with Propidium Iodide and cleared in methyl salicylate. Scales bars true across all images and = 800 µm.

**Supplemental Figure 15 - Concatenated genome improves read alignment percentage for heterografted pepper and tomato** (Supports Figure 3). Each sample, two self-grafts and two heterografts were aligned to the pepper (cvCM334) and tomato (*Solanum lycopersicum* cv Heinz) reference genome. The two heterografts were also aligned to a reference genome that concatenates the pepper and tomato reference genome (referred to as concatenated genome). PP = pepper:pepper self-graft, TT = tomato:tomato self-graft, PT = pepper:tomato heterograft, TP = tomato:pepper heterograft.

**Supplemental Figure 16 - Principal Component analysis (PCA) of the RNAseq samples** (Supports Figure 3). FPKM values of the tomato genes (A) and pepper genes (B) were used to perform a PCA analysis. In the PCA plot, each dot represents an RNAseq sample. The samples are plotted in two dimensions using their projections onto the first two principal components. PP = pepper:pepper self-graft, TT = tomato:tomato self-graft, PT = pepper:tomato heterograft, TP = tomato:pepper heterograft, PC = principal component.

**Supplemental Videos 1: Bend test of self-grafted and heterografted stems 30-days after grafting** (Support Figure 1). Representative self-grafted tomato (A) and pepper (B), and heterografted tomato:pepper (C), pepper:tomato (D) plants that were bent from the rootstock and scion to test graft junction integrity.

## Author’s Note

The author responsible for distribution of materials integral to the findings presented in this article in accordance with the policy described in the Instructions for Authors (www.plantcell.org) is: Margaret H. Frank (.

